# Sources of PCR-induced distortions in high-throughput sequencing datasets

**DOI:** 10.1101/008375

**Authors:** Justus M Kebschull, Anthony M Zador

## Abstract

PCR permits the exponential and sequence-specific amplification of DNA, even from minute starting quantities. PCR is a fundamental step in preparing DNA samples for high-throughput sequencing. However, there are errors associated with PCR-mediated amplification. Here we examine the effects of four important sources of error — bias, stochasticity, template switches and polymerase errors — on sequence representation in low-input next-generation sequencing libraries. We designed a pool of diverse PCR amplicons with a defined structure, and then used Illumina sequencing to search for signatures of each process. We further developed quantitative models for each process, and compared predictions of these models to our experimental data. We find that PCR stochasticity is the major force skewing sequence representation after amplification of a pool of unique DNA amplicons. Polymerase errors become very common in later cycles of PCR but have little impact on the overall sequence distribution as they are confined to small copy numbers. PCR template switches are rare and confined to low copy numbers. Our results provide a theoretical basis for removing distortions from high-throughput sequencing data. In addition, our findings on PCR stochasticity will have particular relevance to quantification of results from single cell sequencing, in which sequences are represented by only one or a few molecules.

## Introduction

DNA sequencing technologies have improved rapidly during the last two decades. Due to decreased cost and increased speed, DNA sequencing is now a standard technique in molecular biology both for sequence determination and quantification.

Before a DNA sample can be sequenced, a sequencing library must be prepared from the sample. Although the steps in library preparation vary, the protocol almost always involves PCR amplification. However, PCR is imperfect; it introduces both skews and new hybrid or erroneous sequences into the pool of amplified DNA molecules. Our goal here is to obtain a quantitative understanding of various artefacts introduced by PCR.

Here, using pools of carefully designed amplicons and Illumina sequencing, we theoretically and experimentally investigate four processes known to cause sequence misrepresentation after PCR amplification. First, we studied the effects of PCR bias focussing on variable PCR amplification efficiencies as a function of the GC content of individual sequences. GC bias has often been considered a major source of sequence misrepresentation after PCR. It has carefully been measured in high-throughput sequencing data, and suggestions have been made as how to minimize GC bias [1–3]. Second, we studied the stochasticity with which each DNA molecule is amplified at each cycle of PCR. A large body of theoretical work has focused on the stochastic nature of PCR [4–8], mostly concerning its implications for quantitative PCR [9]. Few experiments have addressed stochasticity in PCR amplification, however, and we know of no work carefully considering the impact of the stochastic nature of PCR in sequencing data. Third, we studied template switching, a process by which two templates combine to form a novel chimeric product during amplification [10]. Template switching has previously received attention in the metagenomics community, and tools have been developed to detect and remove such chimeric sequences from pyrosequencing data [11]. While elegant, these tools do not provide a quantitative model of how and when template switches occur, and therefore cannot inform future experimental designs. Finally, we studied polymerase errors and their impact on sequencing results. Again, polymerase errors have long been recognized as important, and tools have been developed to remove erroneous sequences from high throughput sequencing datasets; but little is known about the relative magnitude of polymerase errors compared with other unintended effects of PCR [12–14].

For each of the four processes considered in this work, we formulated a mathematical framework, looked for signatures of the process in sequencing data, and compared our theoretical predictions with the experimental data. Our main conclusion is that PCR stochasticity is the most significant source of skewed sequence representation in our low-input, high-throughput sequencing datasets. Polymerase errors are the next most important source of error. However, erroneous sequences are limited to small copy numbers, and thus have only a small effect on overall sequence representation. GC bias and template switches have only minor effects on sequence representation after amplification.

## Materials and Methods

### DNA oligos and PCR

We ordered four *ultramers* from Integrated DNA Technologies: three different types of barcode pairs (BC1-BC1, BC2-BC2 and BC3-BC3) and an adapter oligonucleotide (Table 1). BC1-BC1, BC2-BC2 and BC3-BC3 contain the Illumina P5-SBS3T sequence followed by a 20 nucleotide barcode, the PhiC31 phage Attachment site L (AttL) sequence and another 20 nucleotide barcode (Fig 1 A). Whereas the barcodes of BC1-BC1 and BC2-BC2 have a balanced base composition, BC3-BC3 has GC rich barcodes with an expected GC content of 80%. The adapter oligonucleotide is 5*′* phosphorylated and contains a 15 nucleotide barcode. This barcode acts as a varietal tag [15] and is used to count the absolute copy number of input sequences. The 15 nucleotide barcode is followed by the reverse complement of the Illumina P7-SBS8 sequence. The 3*′* end of the adapter oligo is phosphorylated to avoid circularization.

**Table 1.**
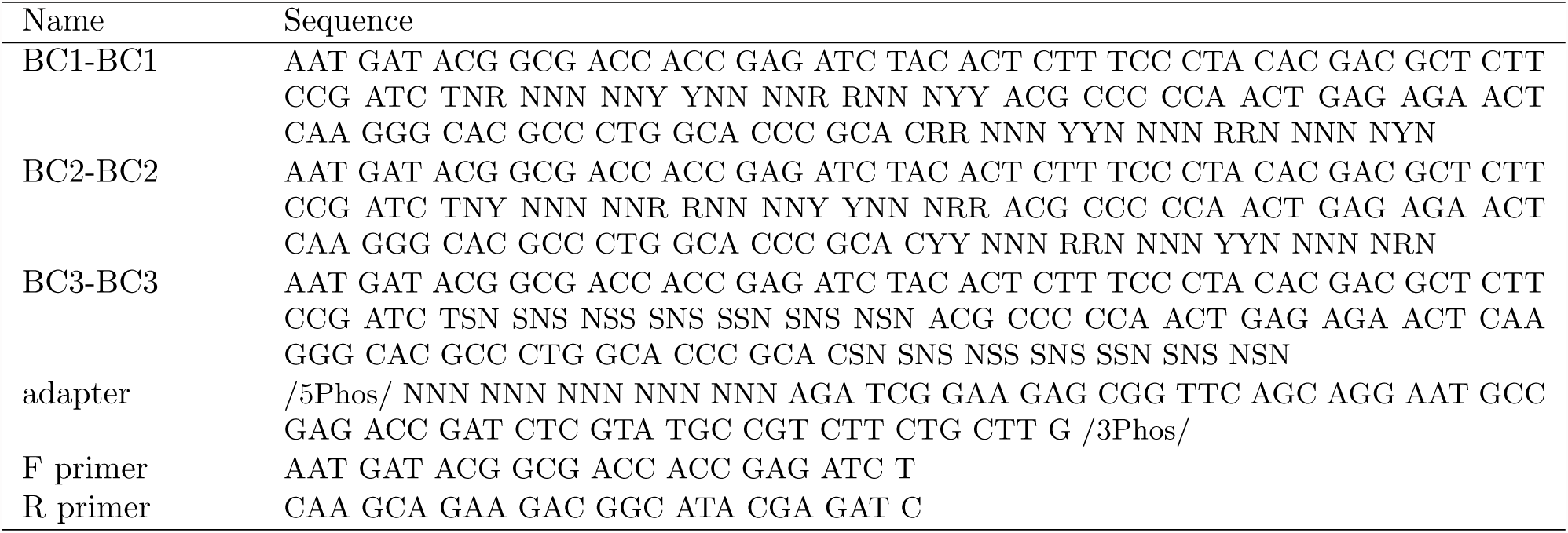
DNA oligonucleotides used

**Figure 1.**
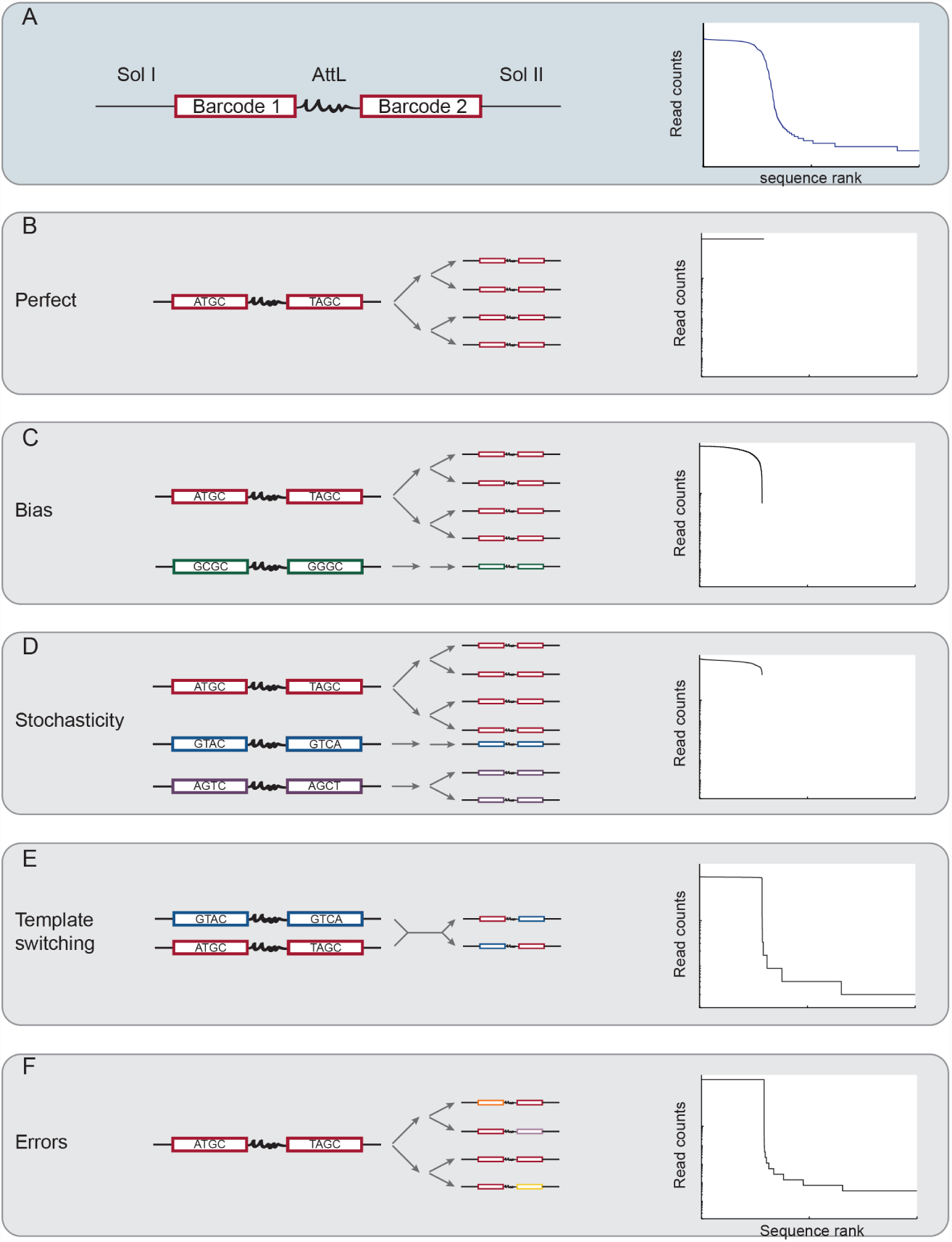
Errors and biases in PCR and their theoretical impact on sequence representation. (A) Left: structure of the amplicons used in this study. Two 20nt barcodes flank a constant sequence (AttL), forming a barcode pair. They are in turn flanked by Illumina P5 and P7 sites as well as sequencing primers (SolI and SolII). Right: sequence rank plot of experimental dataset ZL037. A plateau, a broad shoulder and a long tail (partially visible) are apparent. Schematic representation of perfect (B) and different modes of skewed forms of PCR as well as their expected impact on sequencing data. PCR bias (C) and PCR stochasticity (D) skew the relative abundance of input sequences, but do not add any new sequences to the dataset. In contrast, PCR template switching (E) and polymerase errors (F) generate novel sequences.

We pooled BC1-BC1 and BC2-BC2 in equal amounts and ligated them to the 3’ adapter using CircLigase I ssDNA ligase (epicentre; previously sold as Thermophage ssDNA ligase) as previously described [16]. For the high GC dataset ZL053 we additionally included BC3-BC3 in the ligation reaction at a fivefold reduced concentration relative to BC1-BC1 and BC2-BC2. The ligation reaction was cleaned up with Agencourt RNAClean XP beads (Beckman Coulter) according to the manufacturers instructions. The ligated products were subjected to 25 cycles of PCR using 47*μl* Accuprime Pfx SuperMix (Invitrogen), 1*μl* of 10*μM* forward and reverse primers each (Table 1) and 1*μl* input. Cycling was performed in a BioRad MyCycler Thermal Cycler using standard Accuprime protocol with 58^°^*C* annealing temperature and 30 seconds extension time. The PCR product was gel extracted and sequenced on a single lane of a HiSeq 2000 machine at PE101 per dataset.

### Data processing

Illumina sequencing resulted in 60, 215 and 224 million reads passing filter for datasets ZL037, ZL052 and ZL053 respectively. We merged the paired end reads into their consensus sequence with the Pear tool [17] using standard settings and requiring a consensus sequence of 101 nt. We trimmed and preprocessed the remaining consensus reads using Matlab requiring a perfect match to the constant AttL region, and used the remaining sequences for all subsequent analysis (Table 2) (*preprocessing.m*). All original data files are freely accessible on the Sequence Read Archive under accession SRP057767.

**Table 2.** Sequencing datasets

| Name | BC1-BC1 | BC2-BC2 | BC3-BC3 | reads pass filter | PEAR consensus reads |
| --- | --- | --- | --- | --- | --- |
| ZL037 | + | + | — | $60 * 10^6$ | $15.7 * 10^6$ |
| ZL052 | + | + | — | $215 * 10^6$ | $21.2 * 10^6$ |
| ZL053 | + | + | + | $224 * 10^6$ | $19.6 * 10^6$ |

### Data scaling

The three datasets used in this study differ slightly in the number of input sequences and in sequencing depth. To make direct comparison easier, we linearly scaled the x and y dimensions of ZL052 and ZL053 to match ZL037 by minimizing the squared error between them. Scaling coefficients are reported in Table 3. All further analysis was undertaken on the scaled datasets.

**Table 3.**
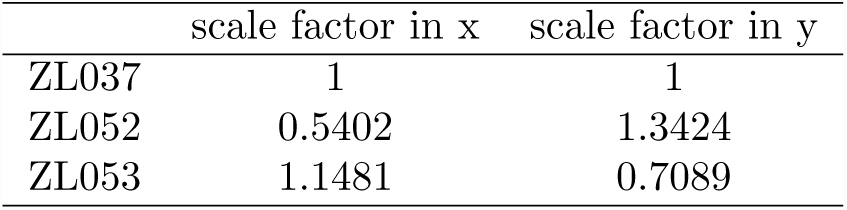
Scale factors

**Table 4.** Variables

| variable | section first used | meaning |
| --- | --- | --- |
| $j$ | Perfect PCR | PCR cycles |
| $N_0$ | Perfect PCR | initial copy number |
| $n(j)$ | Perfect PCR | number of molecules after $j$ cycles |
| $N_{0,x}$ | PCR bias | initial copy number of sequence x |
| $n_x(j)$ | PCR bias | number of molecules with sequence x after $j$ cycles |
| $c_x$ | PCR bias | PCR efficiency of sequence x |
| $m_x$ | PCR bias | estimated PCR efficiency of sequence x |
| $P_{\text{amp}}$ | Stochastic amplification | probability of amplification per cycle |
| $s_0$ | Template switching | per molecule probability of switching |
| $s_j$ | Template switching | probability of template switching on cycle $j$ |
| $S_j$ | Template switching | total number of template switched on cycle $j$ |
| $Q_m$ | Template switching | total number of template switched molecules after $m$ cycles |
| $N_k$ | Template switching | total number of molecules after $k$ cycles |
| $J_k$ | Template switching | approximation of $\frac{Q_k}{N_k}$ from the data |
| $c$ | Polymerase errors | PCR efficiency |
| $R_j$ | Polymerase errors | number of faithfully copied molecules after $j$ cycles |
| $e$ | Polymerase errors | probability of making one error per molecule per cycle |
| $Z_k$ | Polymerase errors | total number of molecules containing a single error after $k$ cycles |
| $F_k$ | Polymerase errors | fraction of all sequences containing errors |

### GC bias

PCR efficiencies were determined as a function of GC content as described in the text. For simulation of GC biased PCR, we calculated the average PCR efficiency for bins of 0.05 width, and normalized to a PCR efficiency of 1.9 for the bin 0.5 to 0.55. We then randomly sampled a binomial distribution to obtain the GC content for 2900 input sequences, and assigned to each sequence the PCR efficiency of its corresponding bin. All sequences were amplified for 25 cycles, and the resulting molecule numbers poisson sampled to simulate sequencing. We chose *λ* as to match the sum of all the reads in the plateau of ZL037.

### Calculation of the probability distribution function of copy numbers after PCR

In PCR, the number of offspring molecules of every sequence in every amplification cycle is drawn from a binomial distribution. When molecule numbers are small, the binomial distribution is poorly approximated by a Gaussian distribution. It is therefore essential to explicitly calculate the binomial distribution of offspring molecules in the early cycles of PCR. However, later in PCR, when molecule numbers exceed around 20 copies, the binomial distribution is efficiently approximated by a Gaussian distribution (Law of large numbers). In our analysis we exactly calculated the Probability Distribution Function (PDF) of copy numbers after PCR for the first 15 cycles, and then switch to a Gaussian approximation for computational ease.

To obtain the exact PDF for the first 15 cycles of PCR, we proceeded as follows. Let us use the vector **S** to denote the distribution of copies of a given barcode-pair after *j* cycles. Each element **S**(*i*) of **S** is the probability of having *i* –1 copies. Thus, **S**(1) is the probability of having 0 copies, **S**(2) of having 1 copy, etc. If eg **S**(3) = 1, it means the probability distribution is a delta function at exactly two copies; if **S**(2) = 0.5 and **S**(3) = 0.5, it means there is a 50-50 chance of having two or three copies. (Note that the largest nonzero element of **S** must in general be *≤* 2^*j*^, so the length of **S** must be *≤* 2^*j+1*^.)

Our approach is to find an updating matrix **M** such that the distribution of copies **S’** on the next cycle is given by **S′** = **MS**.

This formulation exploits the Markovian nature of the process, namely it does not matter how we ended up with k copies on a given cycle; all that matters is we have k copies. To determine the elements of **M**, we first consider an easier problem. Suppose there are k copies on a given cycle; what is the expected distribution of copies on the next cycle? The number of new copies is given by a binomial distribution with parameters k and P; the distribution of total number of copies is k + the number of new copies. Thus, for cases of the distribution **S** where **S**(*i*) is a delta function (S(i)=1), we have *B*(*i, P*) = **MS**(*i*) where B(i,P) is a binomial shifted by i. This means that the *i*^*th*^ column of **M** is B(i,P).

The calculation of the PDF after 15 cycles can be found in *exactpdf.m*.

To approximate the PDF after more than 15 cycles, we used the exact PDF after 15 cycles as a starting point. For position *n* of the PDF after *j -* 1 cycles, we generated a Gaussian *G*_*n*_ with a mean of *n* (1 + *P*_amp_) and a standard deviation of 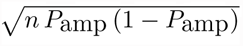. To generate the PDF of copy numbers after *j* cycles, we calculated

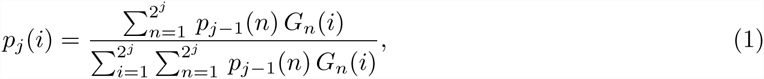

where *i* is the number of copies after and *p*_*j*_(*i*) is the probability of having *i* molecules after *j* cycles. We then iterated this step as required. Code for generating the approximate PDF can be found in *approxpdf.m*.

To simulate stochastic PCR, we randomly sampled 2900 times from the approximate PDF after 25 cycles to obtain molecule numbers for 2900 input sequences after 25 cycles of stochastic PCR. We then sampled from these molecule numbers to simulate sequencing as above.

### Position of template switched reads

We identified half of all template switched reads by comparing dinucleotide anchor sequences between the two barcodes in each barcode pair. We only considered reads where every dinucleotide anchor was either RR or YY, and then checked for barcodes of type 1 joined to barcodes of type 2 and vice versa (Fig 5 B). Using this method, we could identify switches only between barcode classes (i.e. BC1-BC2 or BC2-BC1 type chimeras) and not within them. In our estimate of the per molecule probability of template switching *s*_0_, we assumed that between and within class template switches are equally distributed in the sequence profile. We calculated *J*_*k*_ as the ratio of reads from detected template switched sequences to reads from input barcode pairs and derived the per molecule probability of between class switching as described in the text. *s*_0_ is then simply double that probability.

### Position of single nucleotide errors

We approximated the overall rate of single nucleotide errors in two ways: (1) by determining the minimum hamming distance of each sequence in the shoulder and tail to the plateau sequence and (2) by identifying single nucleotide errors in the dinucleotide anchor sequences, where we quantified the occurrence of RY or YR anchors, that by definition (and with the exception of rare oligonucleotide synthesis errors) are not part of the input barcode pool.

As discussed in the text, we can attribute all errors in sequences with a read count *≥* 3 to polymerase errors. At reads counts less than 3, we cannot be sure whether the observed base pair is due to a polymerase error or just an Illumina sequencing error. When estimating the per molecule polymerase error rate *e* from our data, we did not want to include sequencing errors into the estimate of 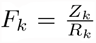, the ratio of reads from erroneous sequences to reads from input sequences. We therefore poisson sampled the sequencing data with *λ* = 0.1, effectively removing all low read sequences and estimated *F*_25_ from the resulting sequence distribution. Such an approximation is valid, as read counts from both erroneous and input sequences are equally affected by sampling, so that *F*_*k*_ remains constant. Indeed, our estimate of the polymerase error rate *e* is robust against different values for *λ* (data not shown). To simulate erroneous PCR on a background of perfect PCR, we simulated perfect PCR, where every molecule present at a given cycle is amplified, and added new sequences caused by errors according to the experimentally derived per molecule error rate, assuming that every new sequence introduced this way is unique. After 25 cycles, we sampled the obtained molecule numbers to simulate sequencing as above.

### Bootstrapping the position of template switched sequences and polymerase errors

To determine whether the distribution of template switched sequences or polymerase errors deviates statistically significantly from a uniform distribution we used a bootstrapping approach. We sampled from the experimental positions of switches or errors with replacement as many times as we found template switches or polymerase errors in the analyzed window. We then calculated the median distance of the drawn positions from the centre of the analyzed window. Repeating this 100000 times, we build up a bootstrapping distribution, and compared the experimentally observed median distance of switches or errors from the centre of the analyzed window to this distribution to calculate the reported p-values.

### Simulation of PCR stochasticity and polymerase errors

We simulated stochastic PCR with polymerase errors using essentially the same model as for simulating erroneous PCR on the background of perfect PCR, but assigned each input and each new sequence a final molecule count sampled from the PDF of molecule counts after 25 − *x* cycles of PCR, where *x* is the PCR cycle at which the sequence first arose. We then sampled these molecule counts as above.

## Results

### Experimental system

To study the effects of PCR on sequence representation in Illumina libraries, we designed an experiment in which sequencing library preparation consists of only a single set of PCR on known, but diverse input sequences, thus removing possibly confounding steps like DNA shearing, reverse transcription or adaptor ligation. We synthesized DNA oligonucleotides containing two regions of random sequence, i.e. two *barcodes*, joined by a constant region. These barcode pairs are flanked by two constant sequences that are required for Illumina sequencing, and which also act as PCR primer binding sites during library preparation (Fig 1 A left).

We subjected about 5000 of these oligonucleotides to 25 cycles of PCR, and sequenced the resulting library for two independent replicates (datasets ZL037 and ZL052; Table 2). We chose 25 PCR cycles to avoid the plateau phase of PCR, while still ensuring sufficient PCR product to obtain a high quality sequencing library. We selected AccuPrime Pfx as PCR enzyme for its superior accuracy, specificity and robustness to GC rich sequences [1]. The relatively low number of input oligonucleotides ensures a high sequencing depth of all input sequences, such that sampling effects during sequencing can be ignored. We expect every input molecule to have a unique sequence, as we used very few input molecules and the combinatorial space of all possible barcode sequences is very large (40 random barcode nucleotides; 4^40^ *≈* 10^24^ possible combinations). This implies that every barcode pair is present at equal abundance in the input DNA pool (i.e. single copy) and should therefore be read out at approximately equal read counts in the sequencing results.

We plotted the experimental sequencing read counts for every barcode pair (Fig 1 A right) sorted from the most abundant to least abundant barcode, that is by sequence rank, where rank 1 is the most abundant sequence. For each dataset, we find that the most abundant barcode pairs are present at similar read counts, forming a plateau in the plot. This plateau is followed by a shoulder of barcode pairs with intermediate abundance and a long tail of low abundance barcode pairs.

Given an equal abundance of all sequences before amplification, we would have naively expected only a plateau. The presence of a shoulder and tail in the experimental data suggests that additional mechanisms are at play. Conceptually, there are a variety of potential artifacts that can be introduced by PCR, each of which will have a different impact on sequence representation after amplification (Fig 1 B-F). We will address each of these mechanisms in turn.

### Perfect PCR

We first considered the sequence distribution we should expect given perfect PCR amplification (Fig 1 B). Perfect PCR faithfully amplifies every molecule in the input DNA pool, simply doubling molecules at every cycle. Relative abundances of different sequences will thus be preserved during amplification.

Mathematically, perfect PCR can be summarized as

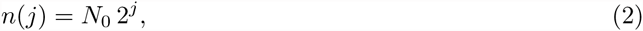

where *n*(*j*) is the number of molecules of a particular amplicon after *j* cycles, and *N*_0_ is the initial copy number of this amplicon.

If every amplicon is unique before amplification, then *N*_0_ = 1, and every sequence will be present at *n*(*j*) = 2^*j*^ copies after *j* cycles of PCR. Plotting copies against sequence rank results in a plateau of height 2^*j*^ reads. If sequencing is deep enough to overcome Poisson sampling effects, we expect a similar plateau when plotting read counts against sequence rank for the Illumina sequencing results, but with a plateau height that reflects sampling during sequencing.

The experimental data deviate substantially from this expectation. A trivial explanation for this disparity is that some sequences were more abundant than others before PCR. We accounted for this possibility by including varietal tags [15] (see Materials and Methods) in our experimental design, which allow absolute quantification in high-throughput sequencing experiments. We were therefore able to count the number of input molecules that carry a particular sequence, and found that all sequences from the plateau were present exactly once before PCR (compare *preprocessing.m*). We therefore conclude that the naive model of perfect PCR does not describe the experimentally observed data.

Below we consider four processes, two of which (PCR bias and PCR stochasticity) introduce skews into the representation of input sequences, and two (template switching and polymerase errors) that generate new sequences during PCR amplification. For easier comparison between the experimental datasets in all subsequent analyses, we normalized the sequence rank plot in x and y to account for differences in input molecule numbers and sequencing depth. The scaled datasets overlap very well (Fig 2; Table 3).

**Figure 2.**
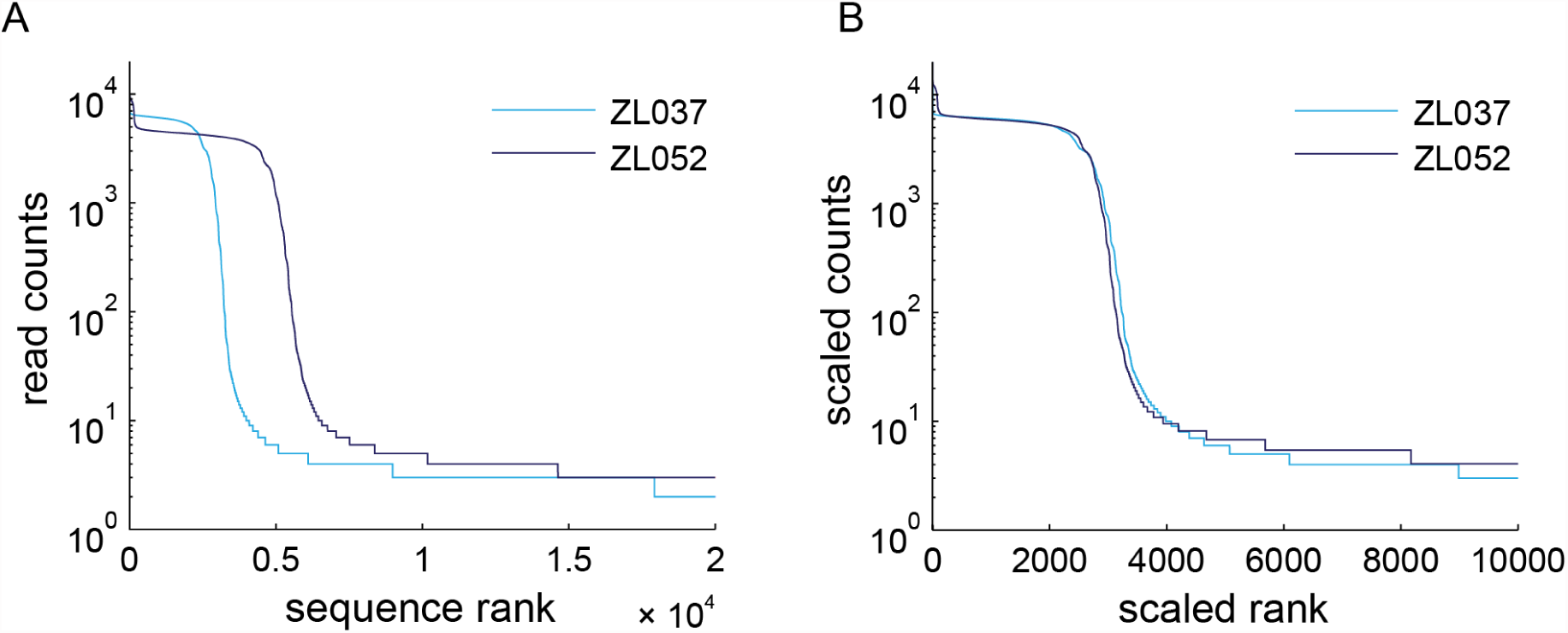
PCR can grossly affect sequence representation in Illumina library generation. Sequence rank plot of replicate datasets ZL037 and ZL052 before (A) and after (B) linear scaling in x and y to compensate for different input amounts and sequencing depth. Scale factors for the x and y dimensions can be found in Table 3.

### PCR bias

The efficiency with which PCR amplifies a sequence may vary from one sequence to the next, depending on factors including sequence composition and secondary structure. A high fraction of G or C can reduce amplification efficiency [1, 3, 18], causing uneven amplification of different sequences in PCR. We therefore tested the possibility that PCR bias was responsible for underrepresented sequences after amplification. Such a bias could contribute to the shoulder or tail in our plots, depending on how strong it was (Fig 1 C).

In equation 2 we assumed perfect amplification on each cycle. GC bias would manifest as a sequence-dependent amplification efficiency. Assuming different amplification efficiencies for different sequences, we can express the expected copy number *E* (*n*_*x*_(*j*)) of sequence *x* after *j* cycles of PCR as

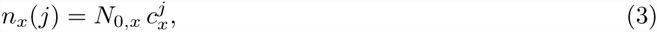

where *N*_0,*x*_ is the initial copy number of sequence *x*, 1 *≤ c*_*x*_ *≤* 2 is its PCR efficiency, and the number of copies *n*_*x*_ is large.

PCR bias, that is a PCR efficiency *c*_*x*_ smaller than the average efficiency of all sequences *c*_*x*_, has been reported in sequences with a high GC content [1]. As GC bias causes uneven amplification of different sequences, we tested whether molecules that were underrepresented after amplification (i.e. the sequences in the shoulder) were rich in GC (Fig 3 A). If GC bias were causing the observed differences in read counts, regions of high read counts should have lower GC contents than regions of low read counts. The experimental distribution of GC content for plateau sequences (high read counts), shoulder sequences (intermediate read counts) and tail sequences (low read counts) are overall very similar to each other (Fig 3 B), suggesting that GC bias is not the primary force shaping the sequence rank curve. We note that the average GC content of plateau, shoulder and tail are statistically significantly different (p-value=1.9 × 10^-4^ using 2-way-ANOVA), but that the effect size is small; plateau sequences have a mean GC content of 0.4463, compared to a mean GC content of 0.4656 and 0.4583 for the shoulder and tail.

**Figure 3.**
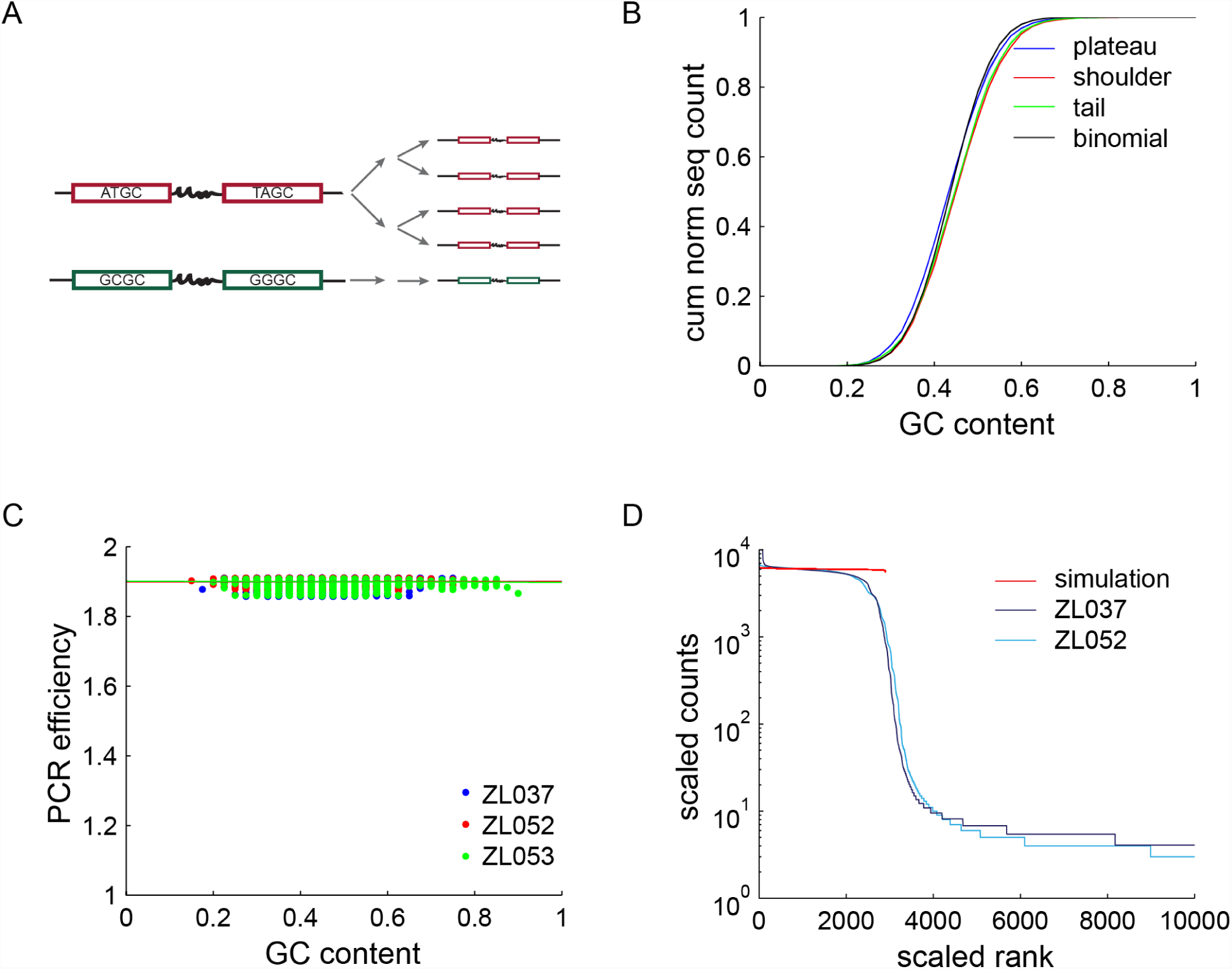
GC Bias. (A) Schematic of two cycles of GC biased PCR. A sequence with balance sequence composition (red) is readily amplified, whereas a GC rich sequence (green) does not amplify well. (B) Cumulative distribution of GC content = / – s.d. for 1500 sequences in plateau, shoulder and tail of the sequence trace of ZL037 and ZL053. No striking differences can be observed. However, the mean GC contents of the three distributions are statistically significantly different. (C) Relative PCR efficiencies as a function of GC content as measured in the plateau of all three datasets, normalized to an efficiency of 1.9 for GC contents of 0.5 to 0.55. Linear fits are plotted as lines. PCR efficiencies are roughly constant across the observed range of GC contents, including the high GC barcode pairs of ZL053 (green). (D) Simulation of PCR with using PCR efficiencies as derived in (C), compared to ZL037 and ZL052. The simulation fails to capture the shape of the data, confirming that GC bias is insufficient to explain the observed sequence distribution.

To investigate if even the observed small differences in GC content can result in sequence misrepresentation, we directly measured the PCR efficiency *c*_*x*_ as a function of GC content in our datasets. Because each input sequence was present as only one molecule before PCR, we could quantify the relative PCR efficiencies of all sequences. Briefly, we determined the GC content of all input sequences, i.e. of all sequences found in the plateau, and calculated their relative abundance. We normalized the relative abundance of each sequence to the mean abundance of all sequences with a GC content between 0.50 and 0.55 to obtain *m*_*x*_, which acts as an estimate for PCR efficiency. Assuming a PCR efficiency of 1.9 for sequences with a GC content between 0.50 and 0.55, we then calculated absolute PCR efficiencies to each GC content bin using the following formula:

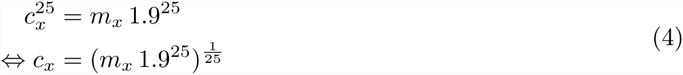

The random design of our barcode sequences naturally leads to few GC rich sequences in the input DNA pool. To improve our ability to calculate *c*_*x*_ at high GC contents, we prepared a third dataset (ZL053), similar to our initial datasets, but into which we spiked about 10% high GC barcode pairs (BC3-BC3) that have an average GC content of 80% (Table 2, Supplemental Fig 1).

Drawing on all three datasets, we find that the PCR efficiency is almost constant across different GC contents (Fig 3 C; linear regression resulted in slopes of 0.0009, 0.0011, and -0.0033 for the three datasets respectively, with 95% confidence intervals *<* 0.008).

Using these empirically measured values for *c*_*x*_, we then simulated PCR and compared the resulting trace to our experimental datasets (Fig 3 D). In agreement with the previous measurements, we find that GC bias as observed in our dataset cannot explain a large amount of the skew observed in the experimental data.

In conclusion, we find only weak evidence for GC bias in our experimental results. This shows that GC bias is not an important force in skewing sequence representation under our experimental conditions. We note, however, that in our experiments the maximum length of a GC biased stretch of nucleotides is *≤* 20nt — the length of a barcode. Longer GC biased regions might well introduce larger biases during PCR. Notwithstanding this caveat, the observed shoulder and tail in our data are unlikely to be formed by GC bias.

### Stochastic amplification of low copy number amplicons

A second source of uneven amplification in PCR is stochasticity. If PCR were perfect, every single molecule would be replicated every cycle. However, PCR is imperfect, so each molecule undergoes replication with a probability of less than 1. For example, if *P*_amp_ = 0.9, then out of every ten molecules amplified per cycle, PCR will fail to replicate one. This is not particularly concerning when PCR is used on DNA mixtures where every sequence is present in high copy numbers. In this case, the expected 1 + 0.9∗1 + 0.1∗0 = 1.9 fold increase of molecule number of per cycle is sufficient to describe the behaviour of PCR.

However, when sequences are present at very low copy numbers, stochastic amplification may have a significant impact on sequence representation. Consider an example. First, consider a *lucky* amplicon that undergoes replication on the first cycle, so that on cycle 2 there are 2 copies. Further suppose that both copies are *lucky* and again undergo replication, so on cycle 3 there are 4 copies (Fig 4 A, red). Compare this to an *unlucky* amplicon (Fig 4 A, blue), which fails to get copied both cycles (for *P*_amp_ = 0.9, this happens (1 – *p*)^2^ = 0.1^2^ = 0.01 or 1% of the time). If their luck evens out and both amplicons get amplified equally during subsequent cycles, the lucky barcode will appear at a copy number of about 4 times more than the unlucky one. This suggests that the distribution of copy numbers for *P*_amp_ = 0.9 will range over more than a factor of 4. Stochasticity in PCR could therefore explain the shoulder observed in the sequencing trace [9].

**Figure 4.**
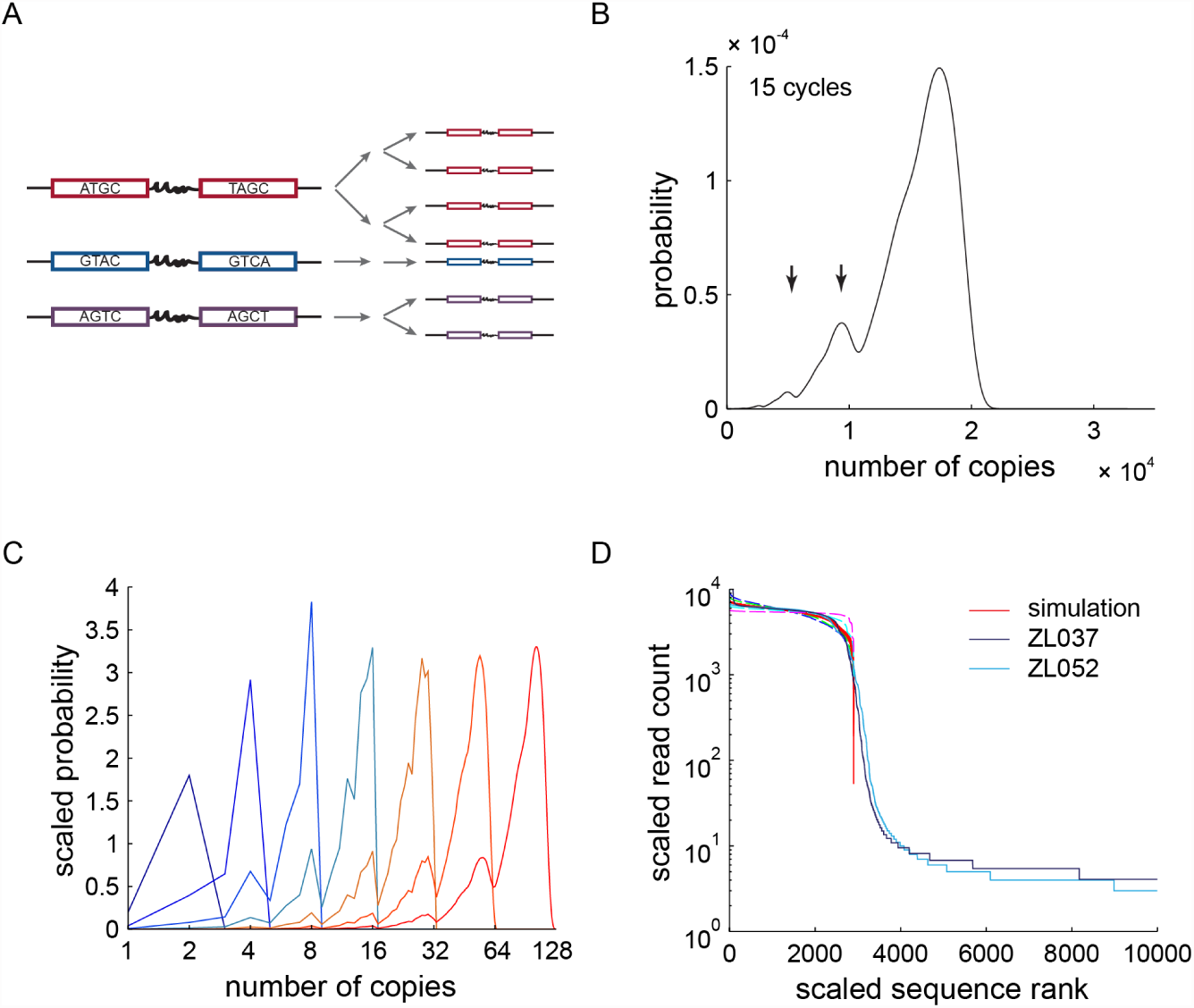
PCR stochasticity. (A) Schematic of two cycles of stochastic PCR amplification. A *lucky* barcode pair (red) gets amplified at every cycle, whereas an *unlucky* barcode pair (blue) fails to get amplified at all. A barcode pair with *mediocre* luck is depicted in purple. (B) The exact probability distribution of sequence copy number after 15 cycles of PCR with *P*_amp_ = 0.9. Arrows indicate two local maxima in the PDF at roughly half and quarter of the molecule numbers as the global maximum. (C) Probability distribution of sequence copy number after 1 to 7 cycles of PCR (blue fading through orange to red). The birth and evolution of the two local maxima observed in (B) is visible. The probability distributions after *j* = 1‥7 cycles were normalized to sum to 2^*j*^ to aid visualization. (D) A sample of 2900 sequences of the approximate probability distribution after 25 cycles of PCR with *P*_amp_ = 0.9 (red) correlates closely with the 2900 most abundant sequence reads of the experimental data. Simulations for *P*_amp_ = 0.8, *P*_amp_ = 0.85, *P*_amp_ = 0.95 and *P*_amp_ = 0.99 are plotted in dashed lines.

We can express this consequence of PCR stochasticity using the recursive expression

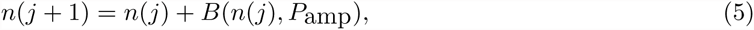

where *B*(*n*(*j*), *P*_amp_) is a binomially distributed random variable with *n*(*j*) trials and *P*_amp_ is the probability of success [5]. This expression is equivalent to modeling PCR as a Galton-Watson process, a stochastic branching process. In this formulation, every node represents one copy of a certain sequence and can give rise to one or two new branches, where each branch corresponds to success or failure of amplification on a given cycle.

Assuming this branching model and a realistic *P*_amp_ = 0.9, we generated the exact PDF of the copy number *n* of a single sequence, starting with a single molecule at cycle 0. After 15 cycles of PCR, the PDF has a clear global maximum (Fig 4 B): As we would expect, most molecules are amplified most of the time. Interestingly, two further local maxima are discernible at copy numbers of 0.5 and 0.25 of the global maximum, corresponding to sequences that missed out on amplification during either one or two of the first two cycles of PCR. The relative heights of these peaks are determined by *P*_amp_.

During the first few cycles of PCR molecule numbers are low, and stochasticity has a large effect. We therefore expected to find the origin of the observed local maxima in the early cycles of PCR. After one cycle of PCR, the PDF is trivial (Fig 4 C). After two cycles, the PDF still shows only one maximum corresponding to molecules amplified in both cycles. After three cycles, the PDF shows two peaks at *n* = 4 and *n* = 8. To reach copy number 8, the molecules have successfully been amplified at every cycle as 2^3^ = 8. To reach *n* = 4, molecules must have failed to replicate during one cycle. The biggest contribution to the probability of *n* = 4 comes from paths where molecules missed out on the first amplification cycle. This fact is immediately obvious when one considers all of the trees giving rise to four branches after three cycles and keeping in mind that a failure to amplify is less likely than success. After 4 cycles, the same structure as observed after 15 cycles becomes apparent. The PDF shows a global maximum at copy number 16 and two local maxima at 0.5 (copy number 8) and 0.25 (copy number 4) of that copy number. When we explicitly calculated the probabilities for these copy numbers, the dominating terms are sequences that failed to amplify on either the first or first two cycles of PCR. A third local maximum is apparent at *n* = 12, which is smoothed out in later cycles. This reasoning confirms that the local maxima in the PDF after 15 cycles correspond primarily to molecules that did not amplify during the first or first two cycles of the PCR reaction.

To test the hypothesis that stochasticity early in the PCR reaction could generate the observed shoulder in the sequence trace, we approximated the PDF of copy numbers after 25 cycles of PCR with a set of constant PCR efficiencies. We sampled from the resulting PDFs to create a profile of read counts vs sequence rank. With a simulated PCR input of 2900 different sequences, we were able to reproduce a shoulder similar to the one observed in experimental data, with the best fit for *P*_amp_ = 0.9 (average correlation coefficient of *R*^2^ = 0.8925 to the three datasets; Fig 4 D). However, the smooth transition of shoulder to tail present in experimental data is missing.

In conclusion, PCR stochasticity has a large impact on sequence representation after PCR amplification and could give rise to most of the experimentally observed shoulder but not the tail.

### Template switching

We next investigated processes producing *new* sequences during library preparation. If a new species is generated during PCR amplification, it will likely be amplified in subsequent PCR cycles like one of the original input sequences. It will, however, lag behind these original sequences by at least one cycle, and will thus be observed less frequently after amplification than most input sequences. Generation of new sequences during PCR could therefore contribute to the shoulder and tail of the sequence trace.

PCR template switching produces hybrid sequences of two sequences already present in the input [10, 19]. DNA polymerase can jump from one template to another in a region of complementarity without aborting the nascent DNA strand during PCR. This nascent strand therefore has a new hybrid sequence, where one piece is complementary to the old template and the other piece is complementary to the new template. Similarly, nascent transcripts can be aborted before completion and then might act as primers in a subsequent cycle of PCR, again resulting in a new hybrid species [10, 19] (Fig 5 A).

**Figure 5.**
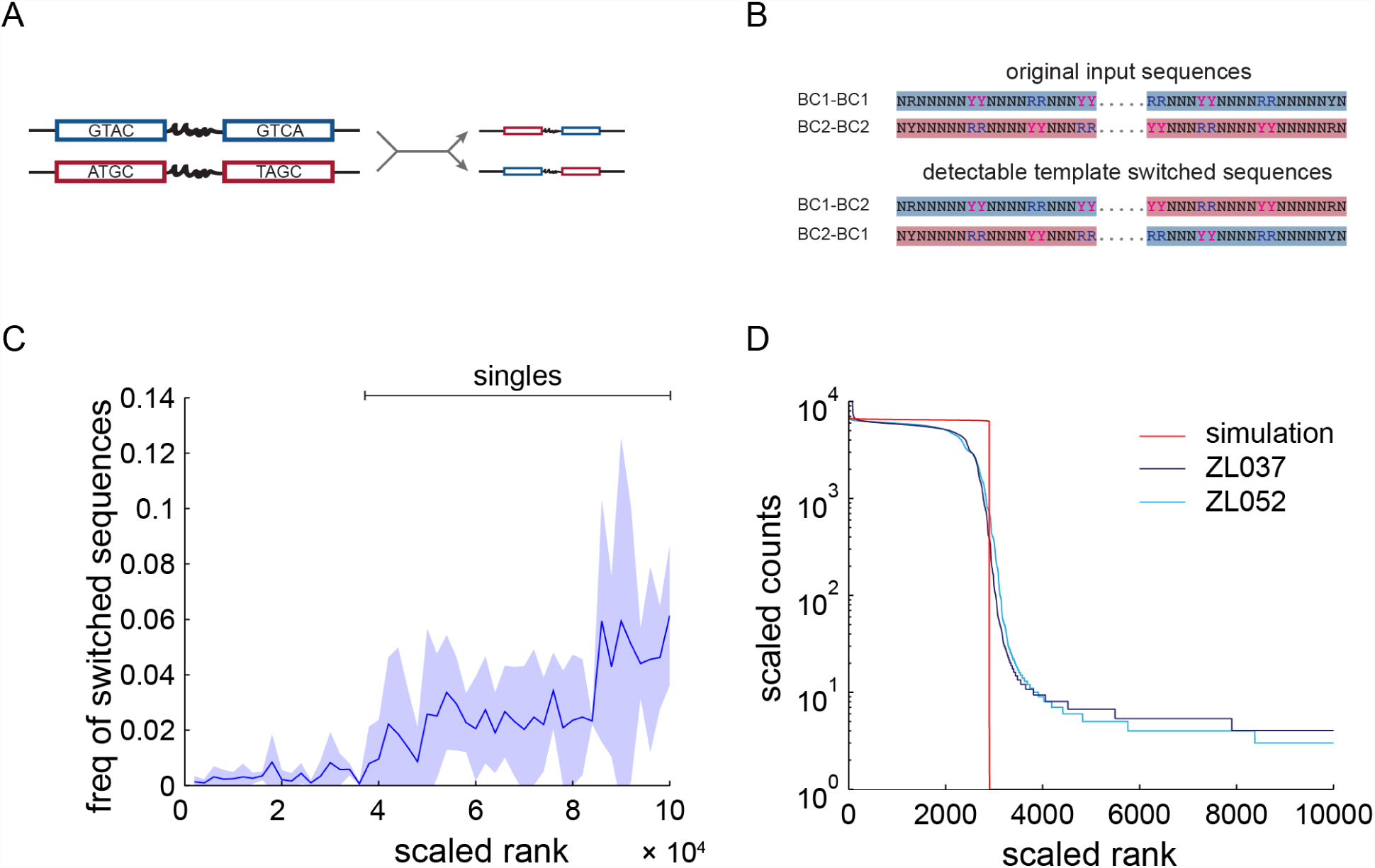
Template switching. (A) Schematic of one cycle of PCR with template switching. During amplification of the blue barcode pair, the polymerase switches to the red barcode pair in the constant region, producing a blue-red chimera. (B) The barcode libraries contain two classes of barcode pairs (BC1-BC1 and BC2-BC2), that are distinguishable by purine and pyrimidine anchors (top). If a BC1-BC2 or BC2-BC1 barcode pair is detected, it must have been formed by a template switch. Such inter-class switches should make up half of all template switches. (C) Abundance of detected template switched sequences +/ – s.d. in sequence rank space. Template switches are rare in abundant sequences, but become more frequent as copy numbers reach one. (D) A simulation of template switching on a background of perfect PCR (red) captures little of the empirical sequence distribution. The only free parameter in our model of template switching, the per molecule rate of template switching *s*_0_, was independently estimated from the data.

To gain a quantitative understanding of template switching, we formulated a mathematical model of the process on a background of otherwise perfect PCR. This model explicitly deals with hybrid sequences produced by polymerase jumping, but mathematically applies equally well to template switching mechanisms where an aborted PCR product serves as an alternative PCR primer. We assume a bimolecular reaction, so that the probability *s*_*j*_ of forming a new product on cycle *j* is governed by the rate of collisions between the two molecules. The collision rate in turn depends on the concentration of the two species, which is proportional to the number *N*_*j*_ of molecules at cycle *j*. Assuming that the probability of template switching following a collision is *s*_0_ and *s*_0_ « 1/*N*_*j*_, then the total number *S*_*j*_ of template switches on cycle *j* is

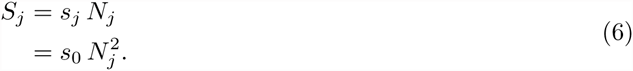

As template switched molecules undergo amplification in every cycle after their generation, the total number of template switched molecules *Q*_*m*_ after *m* cycles is

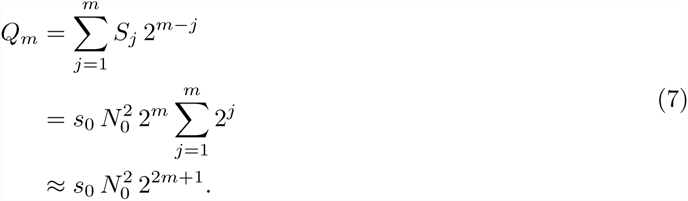

The model predicts that the number of template switches per cycle grows with the square of the total number of molecules in solution *N*_*j*_. As *N*_*j*_ increases exponentially with *j*, the probability of template switching increases exponentially with 2*j*. Accordingly, template switches will become increasingly common in late cycles of PCR, but will not accumulate to levels comparable to the original input sequences, and should be detectable mostly in the tail of the sequence distribution.

To test this prediction experimentally, we searched for signatures of template switching in the sequencing results. Our barcode libraries contained two different classes of barcodes (Fig 5 B). Barcodes of type 1 (BC1) are different from barcodes of type 2 (BC2) at six positions at which the sequence is restricted to either a purine (R=A,G) or a pyrimidine (Y=C,T) base. Based on these anchors, BC1 and BC2 can be reliably distinguished from each other. We started the PCR reaction with a pool of barcode pairs that were either BC1-BC1 pairs or BC2-BC2 pairs for datasets ZL037 and ZL052. Note that BC3-BC3 pairs in the high GC dataset ZL053 are lacking the anchor structure present in BC1 and BC2 and were therefore excluded for this analysis. In the absence of template switching, we would expect to observe only the initial barcode pair types in the sequencing dataset. However, when we detect a BC1-BC2 or BC2-BC1 pair, this barcode pair must have arisen from a template switch across the constant region between the two barcodes. Using this detection method, we are able to detect half of all template switches.

We find that template switched reads are present only at low read counts in the tail of the experimental sequence distribution (Fig 5 C), and that their distribution significantly departs from a uniform distribution (p-values of 0 for all three datasets by bootstrapping). This is in agreement with the our prediction that template switched sequences are created late in PCR. Accordingly, template switches, as detectable by this metric, produce many new sequences (1234, 16190 and 48153 in ZL037, ZL052 and ZL053, respectively), but make up only a small fraction of all reads (8.1×10^-5^, 7.8 × 10^-4^ and 2.6 × 10^-3^). Note that our barcode design favours the the production of chimeras due to the constant region between the two random barcodes.

We then estimated the per molecule probability of template switching *s*_0_ from the data. We define *Q*_*k*_ as the number of template-switched molecules after *k* cycles, which is smaller than the total number *N*_*k*_ of molecules. Although we cannot measure *Q*_*k*_ or *N*_*k*_ directly, we can obtain an estimate of their ratio *J*_*k*_,

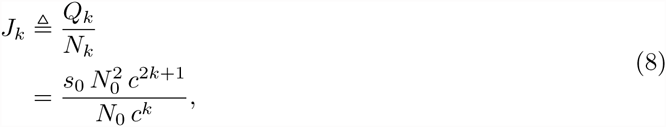

where *c* is the PCR efficiency. We can then solve for the per molecule probability of template switching *s*_0_

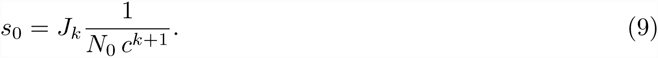

Assuming a PCR efficiency *c* = 1.9, For datasets ZL037, ZL052 and ZL053 after removal of high GC barcodes we obtain a per molecule probability of template switching of *s*_0_ = 2.9 × 10^-15^, *s*_0_ = 1.3 × 10^-15^ and *s*_0_ = 3.3 × 10^-15^ respectively.

Based on the mean value of *s*_0_ = 2.6 × 10^-15^ across our datasets, a simulation of perfect PCR with template switching cannot account for the experimentally observed shoulder (Fig 5 D).

These results indicate that PCR template switching is a rare event in dilute solutions, and only becomes common late in PCR. By then, the newly generated sequences have lost out on many amplification cycles and are present at much lower copy numbers than original sequences. They are therefore detected at low copy numbers in sequencing. Template switched reads do not account for the observed shoulder, and only a small fraction of the sequences in the tail.

### Polymerase errors

A second source of new sequences during PCR are amplification errors. During synthesis of a new DNA strand, DNA polymerase makes errors including single nucleotide substitutions and, at a lower rate, small insertions or deletions [20]. Polymerase error rates strongly depend on experimental conditions, and estimates of error rates vary with the method used to determine polymerase errors. Wild type Taq polymerase is the best studied polymerase used in PCR and is generally used as a relative standard for polymerase fidelity. Estimates of Taq fidelity vary, but are on the order *P* (Error per nucleotide) = 10^-4^ [21, 22]. AccuPrime Pfx polymerase, as used in our experiments, is estimated by the manufacturer to have a fidelity 26× higher than Taq polymerase (*http://tools.lifetechnologies.com/content/sfs/brochures/711-021834%20AccuPrime%20Brochu.pdf*). We therefore expect AccuPrime Pfx to introduce polymerase errors at a probability of roughly

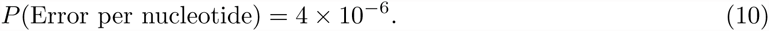

The probability of one or more errors in the 2x20 barcode nucleotides is therefore

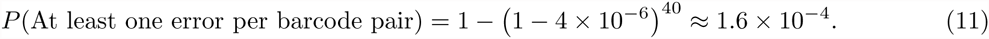

Thus on average, out of every 6250 molecules, a new molecule with at least one error will be produced. These new sequences are subsequently amplified during the remaining cycles of PCR just like any other DNA molecule. However, because, by definition, they lag by at least one cycle of PCR, they are less abundant than original sequences. Moreover, the probability of producing new sequences due to polymerase errors is linearly dependent on the number of amplified molecules, and thus increases exponentially during PCR. Taken together, these considerations suggest that polymerase errors are responsible for part of the shoulder but become increasingly more abundant with low copy number. We therefore predict that the lower part of the shoulder and parts of the tail are formed by polymerase errors (Fig 6 A).

**Figure 6.**
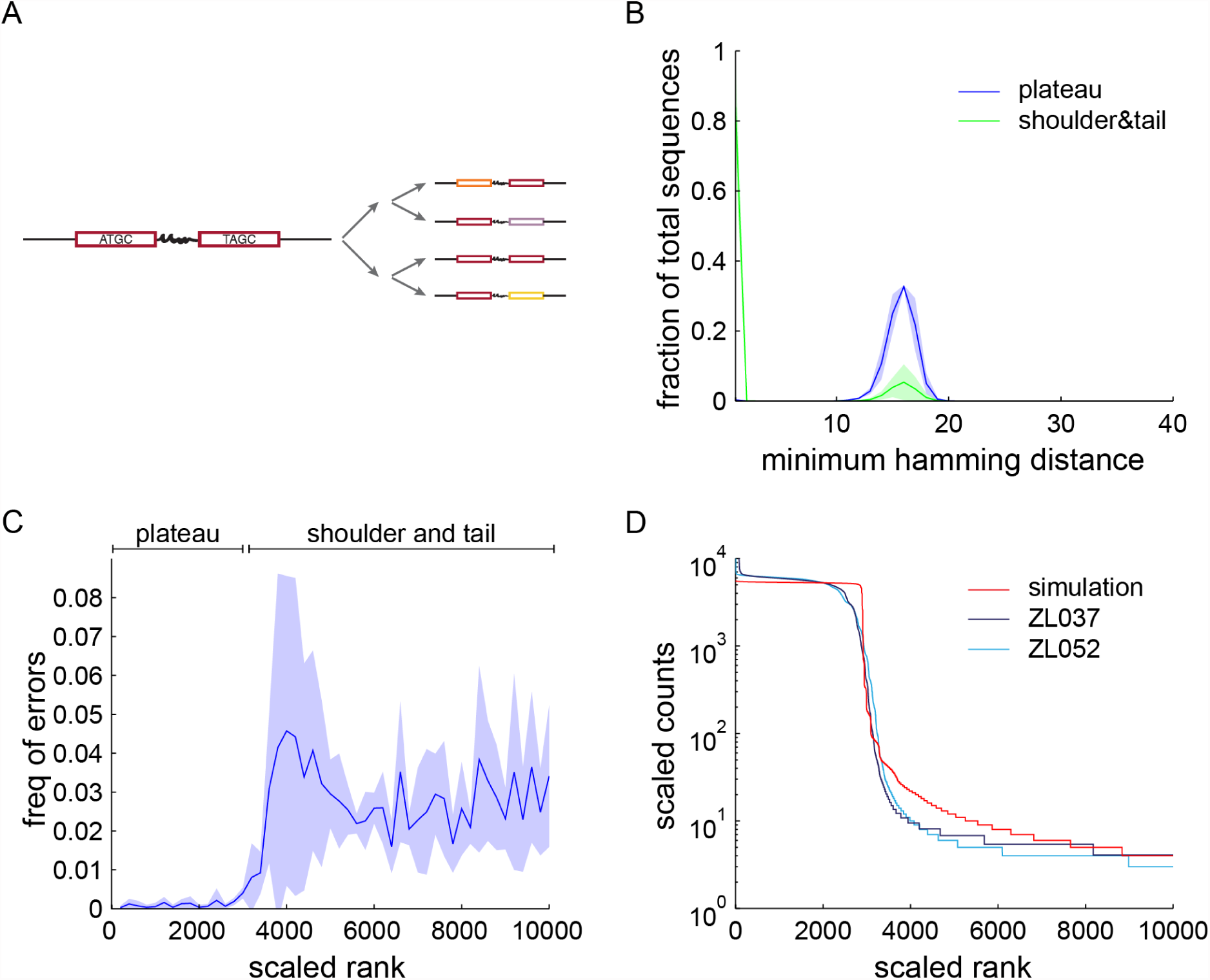
Polymerase errors. (A) Schematic of two cycles of PCR with polymerase errors. Polymerase errors introduce mutations into an input barcode pair (red), effectively producing novel sequences (orange, lavender, yellow). (B) Histogram of the minimum Hamming distance +*/–* s.d. from sequences in the plateau to other plateau sequences (blue) and sequences from shoulder and tail (scaled rank 2900 to 10000) to plateau sequences (green). In contrast to plateau sequences, the majority of sequences from shoulder and tail are within a Hamming distance of one (i.e. one base change) from the parent plateau sequences. (C) Position of errors detected using mismatches to anchor sequences in the barcodes in sequence rank space +*/–* s.d. While the plateau is depleted of polymerase errors, shoulder and tail sequences show a large increase in error frequency. (D) A simulation of polymerase errors on a background of perfect PCR (red) recapitulates the shoulder to tail transition of the observed sequence distribution. The polymerase error rate used for the simulation was independently estimated from the data.

To test this prediction experimentally, we used a Hamming distance metric to identify sequences that arose from polymerase errors. The Hamming distance between two sequences is defined as the number of substitutions necessary to go from one to the other. For example, a sequence with a single PCR error will have a Hamming distance of one to its original parent sequence. Using this metric, we cannot directly differentiate between polymerase errors and Illumina sequencing errors. However, if a sequence appears more than twice in our dataset, it is unlikely that it arose due to an Illumina sequencing error and therefore is either real or result of a template switch or a polymerase error: At a lower bound quality score of *Q*_*phred*_ = 30, that is a base calling error rate of 10^-3^ per nucleotide, the probability of introducing the *same* sequencing error three times into different copies of the same 40nt barcode pair is 40 ∗ 0.001^3^ = 4 × 10^-8^. At a coverage on the order of 10^4^ for *real* sequences — that is, reads from the plateau of the sequence trace — this implies that the probability of introducing the same sequencing error three times into a real barcode pair is 4 × 10^-8^ ∗ 10^4^ = 4 × 10^-4^. With only roughly 5000 real barcode pairs, sequencing errors in sequences present three or more times are negligible. At fewer than three counts per sequence, we cannot exclude the possibility that some of the observed mismatches arise from sequencing errors, especially in the singlet region.

To further reduce the contribution of sequencing errors to our dataset, we sequenced using paired end reads that each span the entire length of the barcode pair, so that every base was sequenced twice. We then determined the consensus sequence of paired end reads using the PEAR tool [17], analyzing only those reads for which a consensus over the whole molecule could be found. Assuming independence of paired end reads, this procedure eliminates the majority of sequencing errors. Taking both read quality and paired end matching arguments into consideration, we will assume that errors identified by the Hamming distance metric arise exclusively from polymerase errors if the sequences have a copy number of more than 2.

We defined the plateau sequences from scaled rank 1 to 2900 as original parent sequences, and find that we can account for about 84% of all other sequences by a single nucleotide substitution in these original sequences. In contrast, the minimum Hamming distances between all the parent sequences are significantly different from each other, such that they could not be related to each other by polymerase errors (Fig 6 B). Due to the length of the tail observed in the data, these findings suggest that the vast majority of all unique sequences in the datasets are actually errors, but all occur at low abundance. Indeed we find that while most of the tail sequences are the product of errors, only roughly 1% (1.1%, 1.6%, 1.5% for ZL037, ZL052 and ZL053, respectively) of all reads derive from errors.

To obtain an independent estimate of the polymerase error rate that does not depend on identifying parent sequences, we quantified polymerase errors by scoring deviations from the expected sequence features of BC1 and BC2 sequences. Each of the defined anchors in BC1 or BC2 is a pair of two purines or two pyrimidines. Any mixed anchor sequence, e.g. AT or CG, therefore must be the result of a substitution. Using these mismatches as measure of polymerase errors, we find that polymerase errors are depleted in the plateau region of the experimental sequence trace (Fig 6 C). The distribution of polymerase errors in this window significantly differs from a uniform distribution as assessed by bootstrapping (p-values=0 for ZL037, ZL052 and ZL053 after removal of high GC spike-ins).

To further quantify the effects of polymerase errors, we estimated the error rate of Accuprime Pfx using our experimental data. Our estimation procedure began by assuming *N*_0_ initial sequences that are amplified with PCR efficiency *c*. After *j* cycles, the total number of faithfully copied molecules *R*_*j*_ is

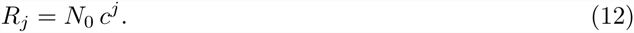

Now let us assume a fixed probability *e* of making an error per molecule on each round. Then the expected number *E*_*j*_ of new errors generated during the *j*^*th*^ round is

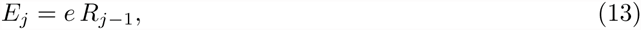

where we use *j* – 1 since the *j*^*th*^ round was produced from *R*_*j*–1_ molecules on the previous round. Erroneous sequences are amplified just like all other sequences. Thus after *k* cycles, there will be *c*^*k-j*^ copies of a erroneous sequence that arose on the *j*^*th*^ cycle. Thus the total number *Z*_*k*_ of molecules containing a single error after k rounds of PCR is

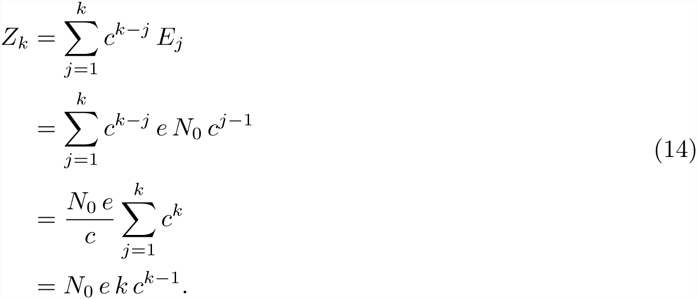

Accordingly, the fraction of all sequences containing errors is

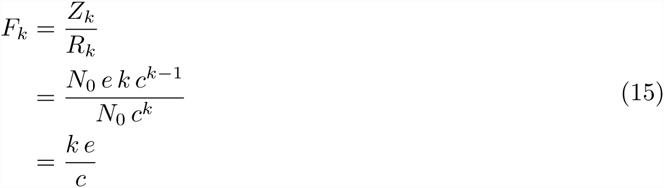

After resampling our sequencing data to remove rare sequencing errors (see Materials and Methods), we can approximate *F*_*k*_ by taking the fraction of all reads in the tail that have a Hamming distance of one to the plateau (the errors) over all reads in the plateau of our data (the original sequences). Therefore, we can solve the above equation for *e* and obtain an estimate for the per molecule error rate in our data. For the three datasets ZL037, ZL052 and ZL053 we obtained values of 0.0008, 0.0012 and 0.0011 respectively. These values correspond to a per nucleotide error rate of 2.0 × 10^-5^, 2.9 × 10^-5^ and 2.8 × 10^-5^, respectively. Note that this experimental polymerase error rate is about 6 times higher than our above estimate based on the manufacturers information and traditional lacZ complementation assays [21, 22].

Based on the relatively steady rate of polymerase errors outside the plateau, we hypothesized that most sequences found at the bottom of the plateau and in the tail of the sequence trace arose from polymerase errors. To test this hypothesis, we simulated erroneous PCR with an overall amplification efficiency of *c* = 1.9 in a deterministic model. Assuming that every individual error is rare, we simulated 25 cycles of PCR on an input of 2900 sequences using the average measured error rate of *P* (Error per nucleotide) = 2.7 × 10^-5^. The resulting sequence rank plot shows a plateau followed by a steep drop off which then softens into a long tail (Fig 6 D). This simulation does not account for the experimentally observed smooth top of the shoulder, but does agree closely with the experimentally observed trace at the bottom of the shoulder.

Taken together, these data confirm our theoretical predictions. Polymerase errors are relatively common, but in absolute numbers happen predominantly late in PCR, and thus are confined to the tail of the sequence distribution, where they make up a large fraction of sequences.

### Stochasticity and polymerase errors explain much of the observed PCR errors

From the above analyses, we expected stochasticity of amplification and polymerase errors to explain most of the observed sequence distribution. To test this hypothesis we simulated PCR by a Galton Watson process, and added polymerase errors at the experimentally observed rate of *P* (Error per nucleotide) = 2.7 × 10^-5^ (Fig 7). Again we assumed that each individual error is rare. Simulation and experimental data show a very good fit, confirming that the observed distribution can be explained by PCR stochasticity and polymerase errors alone.

**Figure 7.**
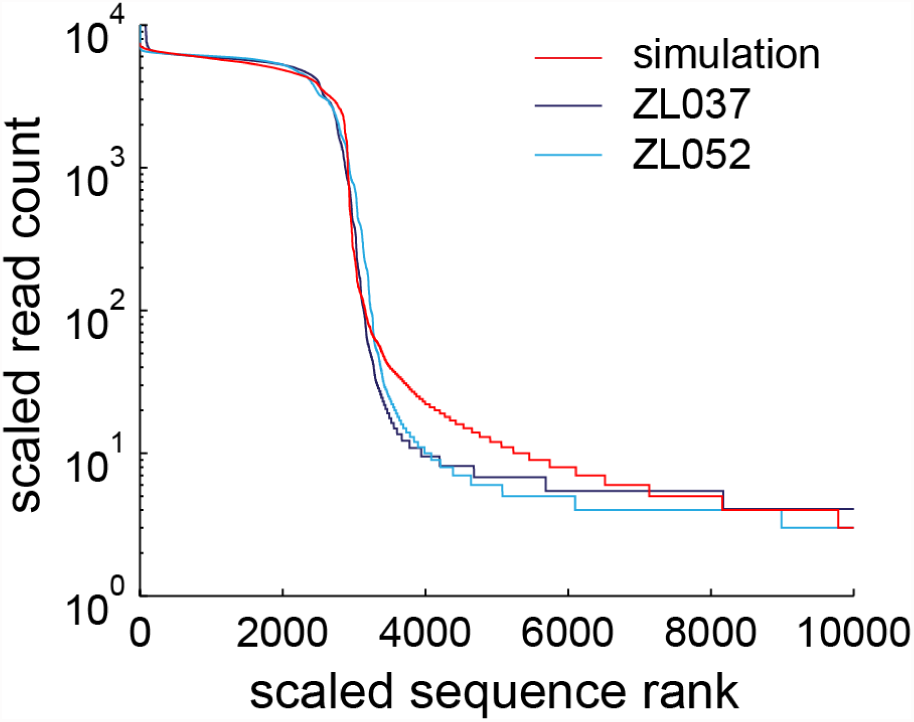
Polymerase errors and stochasticity appear to explain a large fraction of observed data. (A) PCR is simulated as a Galton Watson process with polymerase errors added at the average experimental rate. Simulated (red) and observed sequence profile (light and dark blue) match closely.

## Discussion

We set out to systematically investigate potential sources of sequence misrepresentation in next-generation sequencing libraries, focussing on bias, stochasticity, template switching and errors introduced by PCR. Studying four processes in the same system allowed us to compare their relative importance directly. Using a carefully designed set of amplicons and mathematical models, we find that PCR stochasticity in the first two or three cycles of PCR greatly affects sequence representation of low copy number sequences after amplification. Polymerase errors, and to a lesser extent template switches, generate new sequences which occur predominantly at low read counts, producing the observed tail in the read distribution. GC bias makes only a minor contribution to the observed sequence misrepresentation in the data.

Our findings provide a framework for understanding PCR-induced misrepresentations in sequencing data. Our results on PCR stochasticity have direct relevance for any high-throughput sequencing assay with limited starting material, and are of particular relevance to the single cell sequencing community, where the copy number of target sequences is often equal to one. Our findings emphasize the limits imposed by counting statistics in the low input limit on the quantification of RNA or DNA sequences through sequencing. Stochastic amplification in the first rounds in PCR amplification may contribute to the uneven coverage observed in copy number analyses of single cells [23] or variation of transcript abundance in single cell RNAseq [24, 25]. Indeed, variation of transcript abundance increases with decreasing copy number [24].

To overcome the limits imposed by stochasticity, a number of quantitative tools have been developed to assess and even deconvolve technical and biological noise in single cell RNAseq studies [25, 26]. In parallel to these computational efforts, experimental techniques have been developed to minimize PCR induced distortions in low input sequencing experiments. Techniques like multiple displacement amplification [27], antisense RNA amplification [28] or multiple annealing and looping-based amplification cycles [29] aim to minimize the use of PCR. These approaches exploit linear or quasilinear amplification to minimize the rapid accumulation of errors and biases that arise during exponential PCR amplification and therefore allow for better post-amplification quantification of nucleic acids. In contrast, single molecule barcoding techniques [15] still rely on PCR, but compensate for misrepresentations after sequencing. Individual molecules are uniquely labeled before amplification, so that the number of input molecules can be precisely quantified. Our results underscore the importance of such strategies in cases where copy numbers *<* 8 must be quantitatively resolved.

The introduction of new sequences, through polymerase errors or template switches, are of particular interest in environmental genomic applications where the 16S rRNA gene from a diverse pool of microbes is sequenced to determine the composition of the population. These studies aim to to determine the true number of different 16S rRNA sequences in the dataset, which act as a measure of the number of distinct microbial species in the pool. Algorithms have been developed to remove single nucleotide polymerase and sequencing errors [12, 13], as well as chimeric sequences [11], from high-throughput data. However, because the goal of these algorithms is to identify and remove errant sequences, they do not quantitatively address the distortions in the representation of single molecules that arise from high-throughput sequencing. Our work aims to close this gap, and to provide a quantitative and mechanistic basis for updated experimental designs and analyses. Our data suggest that polymerase errors have a clear signature and, given sufficient sequencing depth, are easily distinguished even from rare input sequences by abundance after amplification. We were further able to measure the critical parameter for a model of template switching, so that we can predict how much we have to dilute our input samples to limit chimeric sequences to an acceptable level.

Although GC bias has previously been reported to introduce misrepresentations into sequencing data [1], we find that it explains little of the observed sequence misrepresentation under our conditions. One reason for this discrepancey may be that effects of GC bias are minimized in our experimental system by the short length of our barcodes. Additionally, we expect that the use of Accuprime polymerase, and of a thermocycler with relatively slow ramp rates (2.5 deg/sec), further diminishes the detrimental effects of GC bias [1].

We designed the experimental system used in this study to disentangle different sources of PCR induced misrepresentation in sequencing datasets. The artificial design of our PCR amplicons allowed us to carefully measure the magnitude of different distorting effects in PCR in a well controlled setting. At the same time, however, this design means that our system differs from many real world applications of high throughput sequencing, so further work will be needed to assess how our findings generalize.

## Acknowledgments

The authors would like to acknowledge Barry Burbach, Ian Peikon and Diana Gizatullina for technical support and Alex Koulakov and Paul Masset for insightful discussions. Additionally, we would like to thank Hassana Oyibo who provided the inspiration for this work.

## Supplemental figures

**Figure S1.**
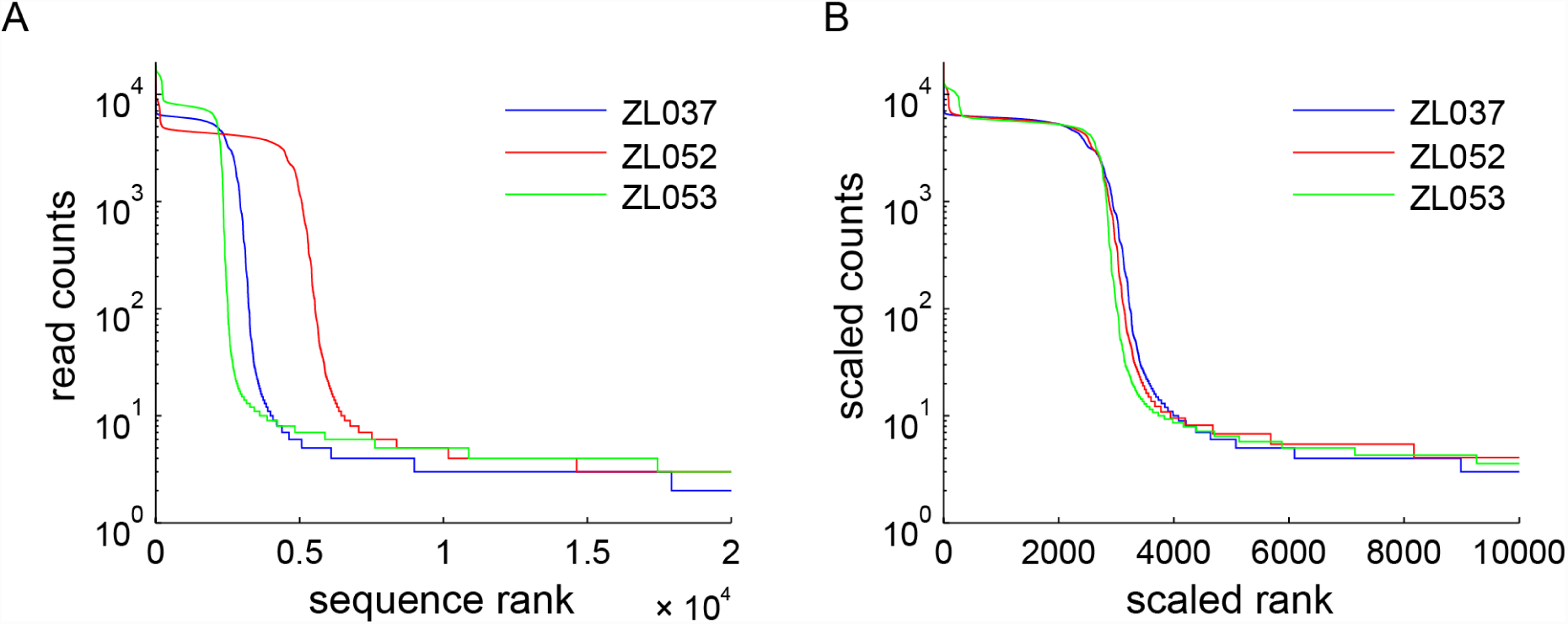
Sequence rank plot of replicate datasets ZL037, ZL052 and high GC dataset ZL053 before (A) and after (B) linear scaling in the x and y to compensate for different input amounts and sequencing depth. Scale factors for the x and y dimensions can be found in Table 3.

